# Novel Cyclic Peptides in Seed of *Annona muricata* are Ribosomally Synthesized

**DOI:** 10.1101/647552

**Authors:** Mark F. Fisher, Jingjing Zhang, Oliver Berkowitz, James Whelan, Joshua S. Mylne

## Abstract

Small, cyclic peptides are reported to have many bioactivities. In bacteria and fungi they can be made by non-ribosomal peptide synthetases, but in plants they are exclusively ribosomal. Cyclic peptides from the *Annona* genus possess cytotoxic and anti-inflammatory activities, but their biosynthesis is unknown. The medicinal soursop plant, *Annona muricata*, contains annomuricatins A (cyclo-PGFVSA) and B (cyclo-PNAWLGT). Here, using *de novo* transcriptomics and tandem mass spectrometry, we identify a suite of short transcripts for precursor proteins for ten validated annomuricatins, nine of which are novel. In their precursors, annomuricatins are preceded by an absolutely conserved Glu and each peptide sequence has a conserved proto-*C*-terminal Pro, revealing parallels with the segetalin orbitides from the seed of *Vaccaria hispanica*, which are processed through ligation by a prolyl oligopeptidase in a transpeptidation reaction.

---

Plants express a number of different types of cyclic peptides. There are several families of homodetic cyclic peptides (i.e. peptides cyclized by linking of their *N*- and *C*-termini) in plants whose biosynthesis has been well-characterized. These are the cyclotides,^1, 2^ cyclic knottins,^3^ and PawS-derived peptides (PDPs),^4,5^ all of which are stabilized by disulfide bridges and are cyclized by a cysteine protease called asparaginyl endopeptidase (AEP).^4,6-9^ As such, these families of cyclic peptides are known as ribosomally-synthesized and post-translationally modified peptides (RiPPs).^10^

There is another group of homodetic cyclic peptides in plants, known as orbitides. These are also RiPPs, but have no disulfide bonds and vary from five to 16 amino acid residues, although most are 7-9 residues.^10,11^ Orbitides are also considered to be RiPPs, but much less is known about their biosynthesis than the peptide families mentioned above. One family of orbitides, the PawL-derived peptides (PLPs), are closely related to the PDPs, and probably cyclized by the same mechanism,^12,13^ but most other orbitides do not contain Asp or Asn residues, which are required for cyclisation by AEP.^10^ Thus most orbitides probably have different mechanisms of cyclisation to PLPs.

The genes encoding orbitides are known only in a small number of cases. Condie et al.^14^ demonstrated that the segetalins, a group of orbitides found in the seeds of *Vaccaria hispanica* (*Saponaria vaccaria* L., Caryophyllaceae), are individually encoded by short genes that express propeptides 30-40 amino acids long. The researchers also showed evidence of a genetic origin for orbitides in some *Citrus* species. Okinyo-Owiti et al.^15^ characterized three novel orbitides from flax (*Linum usitatissimum* L., Linaceae), called cyclolinopeptides 17-19, and found that these cyclolinopeptides (more recently called linusorbs^16^) are derived from gene-encoded precursors, in which multiple peptides are encoded in a single gene. A search of expressed sequence tags from *Jatropha curcas* (Euphorbiaceae) suggested that curcacyclines A and B are genetically encoded.^10^ In these species the cyclic peptide is excised from a highly conserved *N*-terminal leader sequence and *C*-terminal follower sequence (though the *Jatropha* peptides appear to have no follower sequence). The core peptides (the linear precursors of the final cyclic peptide) show little conservation, except at their *N*- and *C*-termini.

There are many other orbitides whose biosyntheses are not known. These include the ∼35 orbitides found in plants of the Annonaceae family, including some reported to possess cytotoxic^17,18^ and anti-inflammatory^19,20^ activities. Two of these orbitides are found in the seeds of *Annona muricata*. Annomuricatin A (cyclo-GPFVSA; monoisotopic mass 558.28 Da) was characterized by Li et al.^21^ and annomuricatin B (cyclo-PNAWLGT) by the same group.^22^ A third orbitide, annomuricatin C, was reported by Wélé et al.^23^ but was later shown by structural studies to be identical to annomuricatin A.^24^

We were interested in the genetic origins of the annomuricatins, which we investigated by combining *de novo* transcriptomics with peptide tandem mass spectrometry. Having sequenced a novel orbitide, annomuricatin D, from tandem mass spectrometry data, we were able to identify a single transcript encoding both annomuricatin D and the previously-known annomuricatin A, thus demonstrating that annomuricatins are ribosomally synthesized. We were then able to identify transcripts encoding a further eight novel annomuricatins (E-L) and sequenced them by mass spectrometry.

## RESULTS AND DISCUSSION

To characterize the biosynthesis of the annomuricatins, total RNA was extracted from *A. muricata* seeds, next generation RNA sequencing (RNA-seq) performed and a transcriptome assembled using established methods.^12^ Using cyclic permutations of the two known peptides annomuricatin A and annomuricatin B, the transcriptome was searched for sequences encoding them; there were hundreds of contigs that had the potential to encode annomuricatin A, but we found none that could encode annomuricatin B.

A liquid chromatography-mass spectrometry (LC-MS) analysis of seed peptide masses for *A. muricata* revealed a mass consistent with the presence of annomuricatin A, no mass for annomuricatin B, but critically it revealed several other seemingly abundant masses (Figure 1). One of these, eluting at 24.7 min, was especially abundant and the protonated molecule had a measured *m/z* of 946.515 [M+H]^+^ (Figure 2A). Treatment of the sample with 1.2 M hydrochloric acid followed by liquid chromatography-tandem mass spectrometry (LC-MS/MS)^25^ caused the formation of a new compound with a molecular weight ∼963 Da, as shown by peaks at *m/z* 482.767 [M+2H]^2+^ and 964.524 [M+H]^+^, although the original peak (now measured at *m/z* 946.514) remained (Figure 2B). If the 946.5 *m/z* ion was a novel annomuricatin, this is consistent with partial acid hydrolysis of its peptide backbone leading to an 18 Da mass increase.^25^

**Figure 1.**
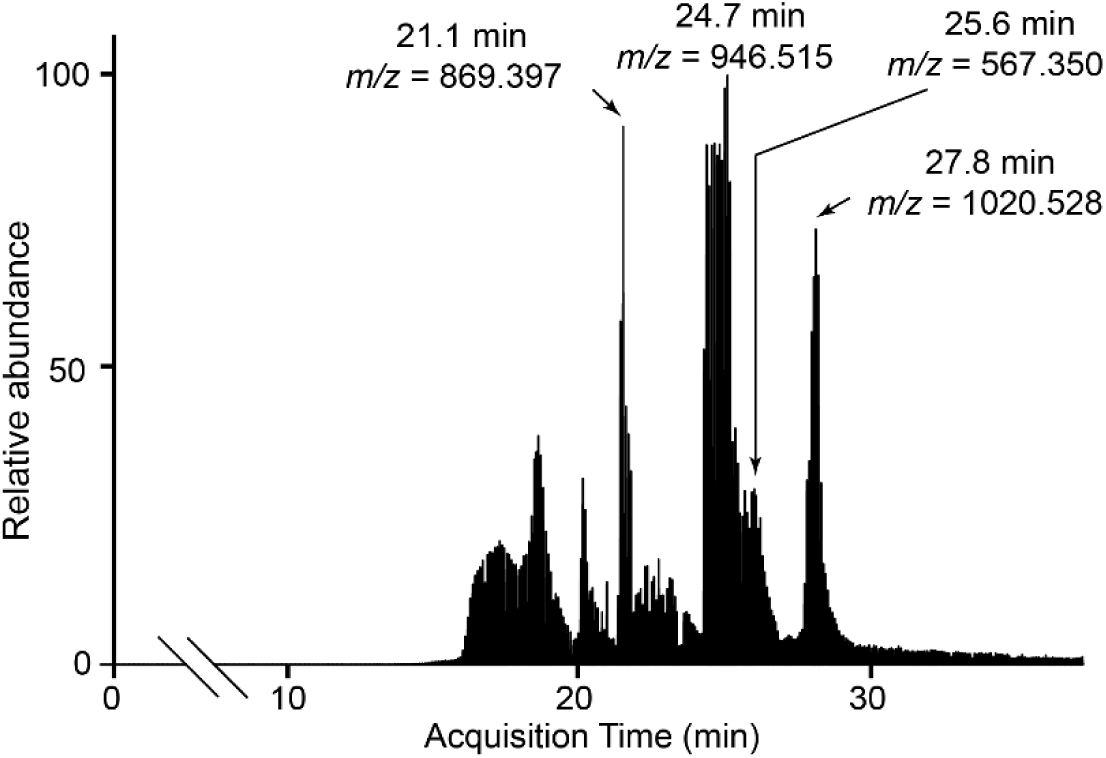
Total ion current chromatogram for extracts of *A. muricata* seeds with the *m/z* values and the acquisition times of the largest peaks labelled. All ions shown are [M+H]^+^.

**Figure 2.**
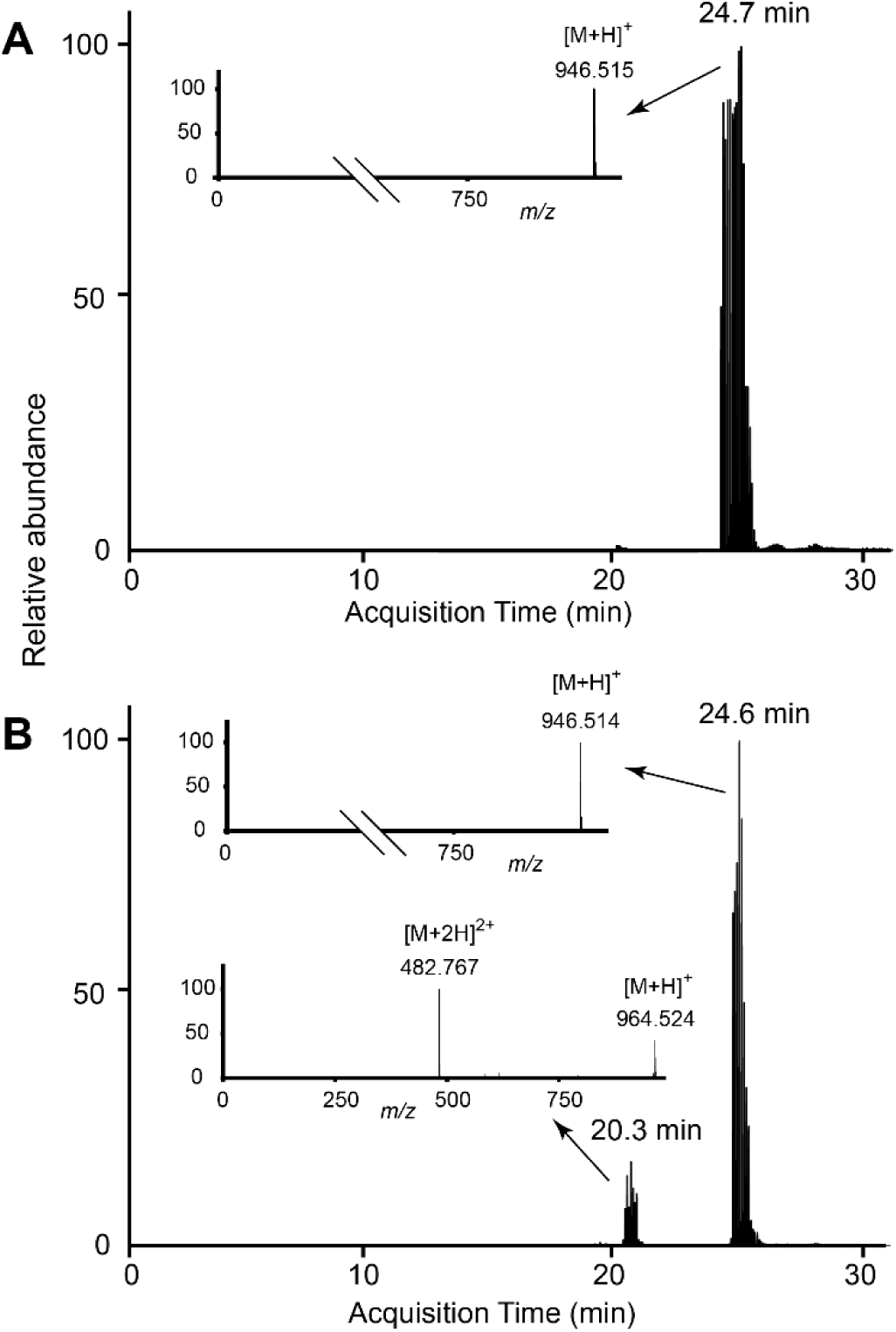
Combined extracted ion chromatograms at *m/z* 946.515 and *m/z* 964.526 for peptide extracts of *A. muricata* seeds. (A) Native extract; (B) After treatment with dilute hydrochloric acid. Insets show the mass spectrum at each peak in the chromatogram.

### Peptide evidence for annomuricatin D

Working on the hypothesis that the *m/z* 946.5 ion was a novel annomuricatin, we sequenced the protonated molecule of the hydrolyzed product (*m/z* 964.5) from MS/MS data to give the sequence SXFPPXPGH (where X represents either of the two isobaric residues Leu or Ile) (Figure 3), corresponding to an exact *m/z* of 964.526 for the acyclic peptide. The order of the Ser and Ile/Leu residues at the *N*-terminus could not be determined unambiguously by MS/MS, however. We tentatively named this novel cyclic peptide annomuricatin D.

**Figure 3.**
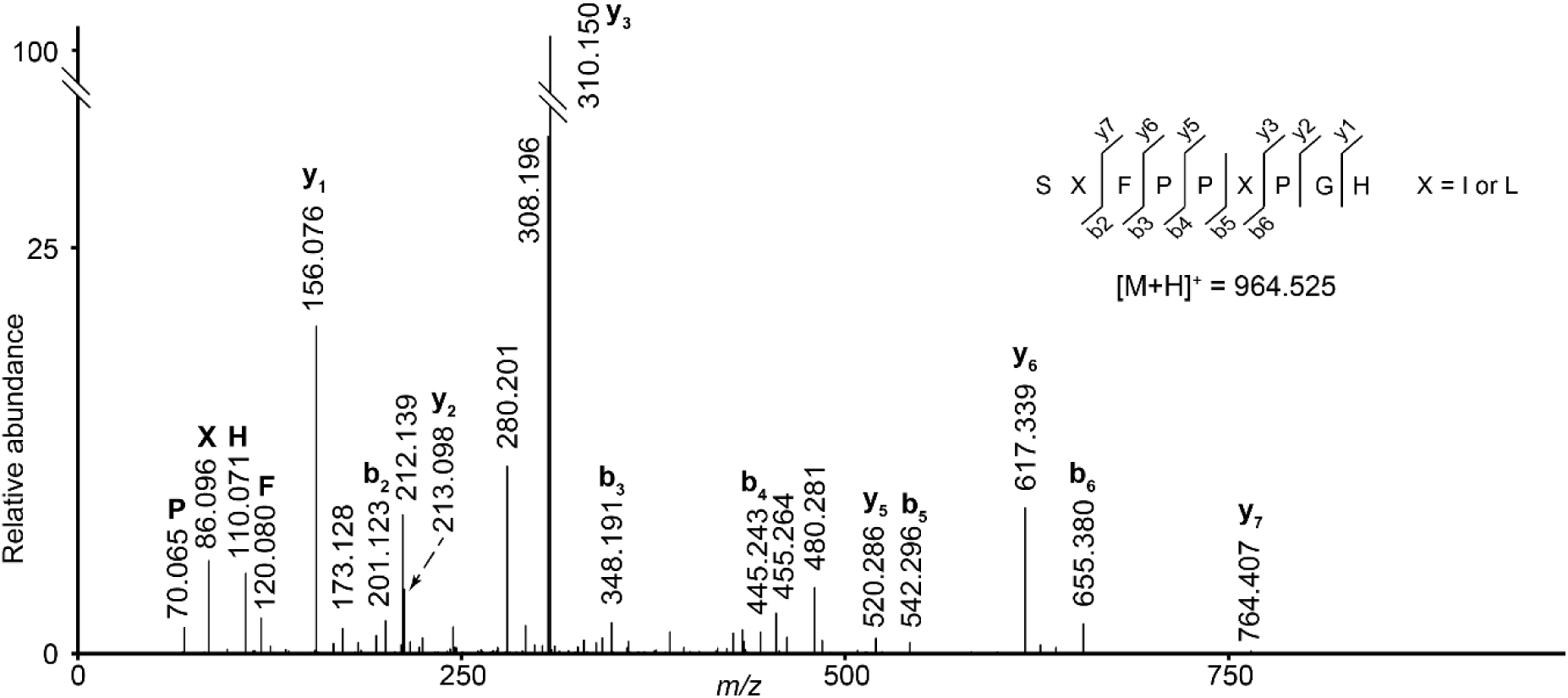
MS/MS confirmation of annomuricatin D. Partial hydrolysis using hydrochloric acid gave a sequence of b and y ions, with the ring breakage *C*-terminal to the His residue (inset shows the peptide sequence). Note that the order of the Ser and Ile/Leu residues could not be unambiguously determined.

### Identification of novel transcripts encoding annomuricatins

Returning to search the *A. muricata* transcriptome using tBLASTn with all 72 possible annomuricatin D query sequences (i.e. all circular permutations of the two possible 9-residue sequences with four possible Ile/Leu combinations) produced just one contig with a perfect match, namely GHSIFPPIP. In the contig matching annomuricatin D, we observed that the complete ORF also encoded annomuricatin A (Figure 4).

**Figure 4.**
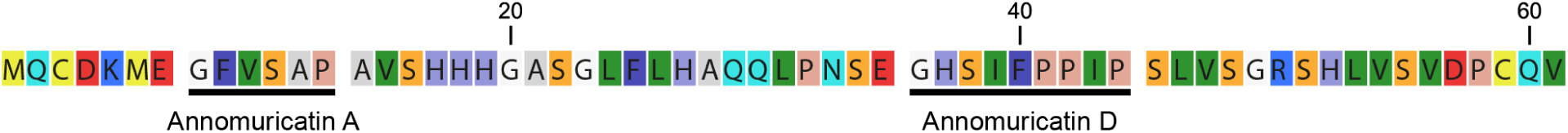
Searches of the *A. muricata* transcriptome by tBLASTn using annomuricatin D as the query sequence revealed a contig whose ORF encoded annomuricatin D as well as the previously published annomuricatin A.

Within this predicted precursor protein, the annomuricatin sequences both ended with Pro and were both preceded by Glu, consistent with the hypothesis that their proteolytic maturation was conserved and involved two different enzymes – one targeting Glu that would release the *N*-terminus of the core peptide and a protease that targeted Pro that would probably perform a cleavage-coupled transpeptidation to the freed *N*-terminus to form a cyclic annomuricatin in a manner similar to other cyclic RiPPs. The sequence for annomuricatin A was *N*-terminal to annomuricatin D in the encoded sequence and so we named the transcript *<u>P</u>ro<u>a</u>nno<u>m</u>uricatinAD*, abbreviated to *PamAD*. Using the *PamAD* transcript as a tBLASTn query, five similar coding sequences with the potential to encode nine annomuricatins were discovered, with a maximum expected value of 4×10^−3^.

Consistent with the lack of peptide mass evidence, no evidence was found for a transcript matching the previously reported annomuricatin B sequence.^22^ This variation in peptide content is something that may be due to environmental differences in transcript expression.^26,27^ To confirm the sequences obtained by RNA-seq and assembly of contigs, primers were designed against the end of each contig to amplify a full length ORF and used genomic DNA as the template. We were able to amplify all six genes and found them to be intronless and a 100% match to the contigs assembled by RNA-seq.

### Confirmation of annomuricatins E to L by tandem mass spectrometry

Knowing the sequence of potential peptides facilitates their confirmation by LC-MS/MS, even without having to linearize the peptides before MS/MS analysis. Using the gene-encoded sequences, we predicted the expected mass for each peptide and could identify masses that matched predictions in all the putative *Pam* genes (Table 1). These putative masses named annomuricatin E to L were each sequenced by LC-MS/MS (Figures S1-S10). Annomuricatins G and I were found in two forms having either a reduced (Figures S3 and S6 respectively) or oxidized methionine (Figures S4 and S7). It is not possible to say whether this is a biologically relevant post-translational modification or occurred during peptide extraction and purification. We have therefore not classified the two forms as separate peptides.

**Table 1.**
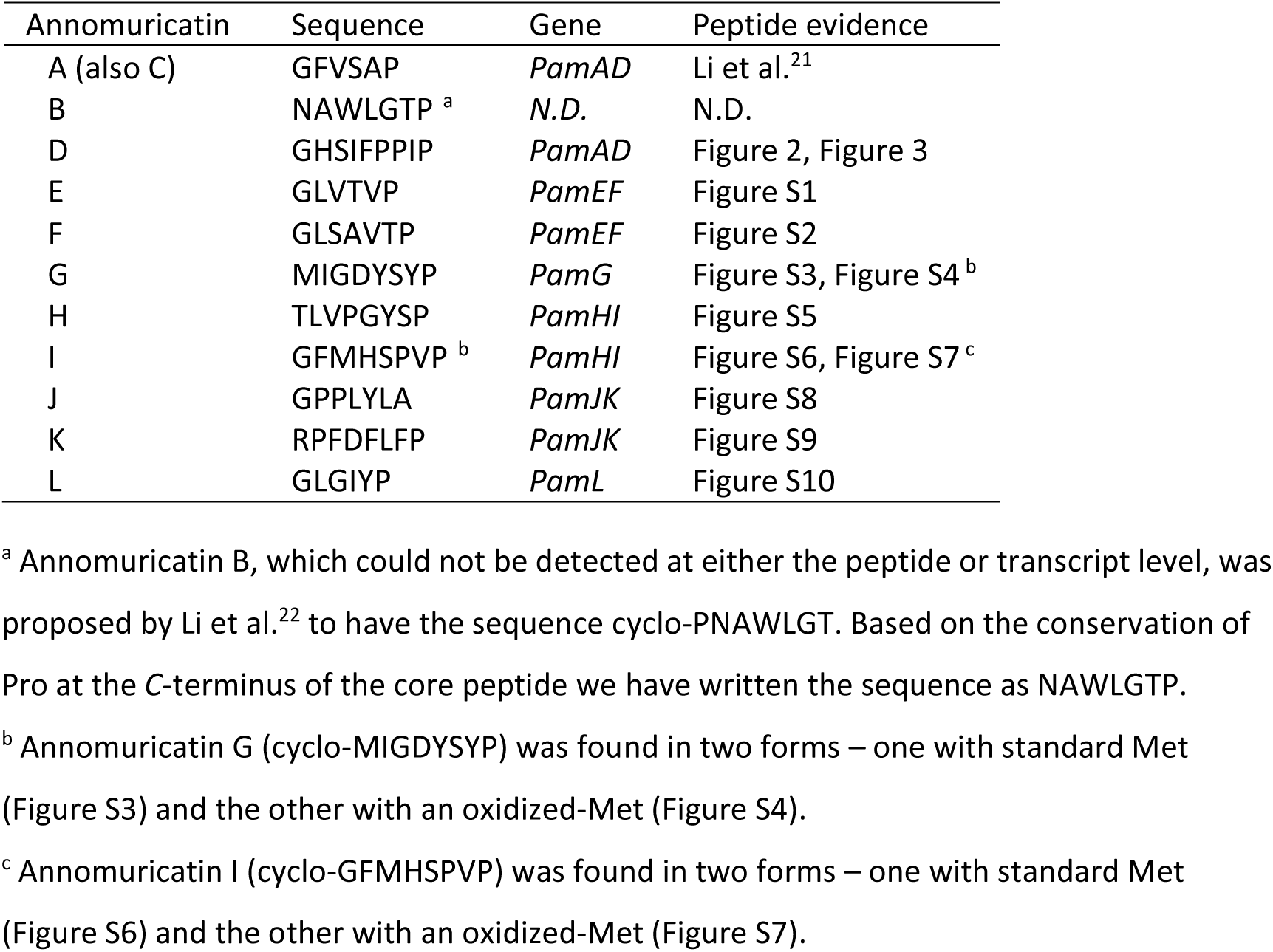
Summary of annomuricatin cyclic peptides, the genes that encode each and evidence for the peptides from mass spectrometry. N.D. = not detected.

Although peaks were found in the mass spectrum corresponding to a second putative annomuricatin in the *PamL* transcript, MS/MS data were not consistent with the predicted sequence from the transcript (Figure S11). We also searched for a possible second peptide in the *PamG* transcript, but none of the sequences searched was found in the mass spectrometry data.

Like many other cyclic peptides, most of the core peptide sequence is not conserved, except for the *N*-terminal and *C*-terminal residues. In the case of the annomuricatins, the *N*-terminus is most often Gly, which tends to be the case in the majority of cyclic RiPPs, and the *C*-terminus residue is Pro or, in one case, Ala, both of which can be cleaved by a prolyl oligopeptidase.^28^ In the leader sequence, Glu in the P1 position to the core peptide is absolutely conserved and the P2 residue varies, but is most often Ser. Assuming that the precursor peptide is cleaved at the proto-*N*-terminus of each of the two core peptides prior to cyclization, as is thought to be the case for the segetalins of *Vaccaria hispanica*,^*29*^ then the only other very highly conserved residue is found at the fourth residue from the *C*-termini of the two propeptides thus formed. This residue is invariably Pro (Figure 5), and it is tempting to speculate, despite its distance from the core peptide, that this is in some way required for the prolyl oligopeptidase to cyclize the peptide.

**Figure 5.**
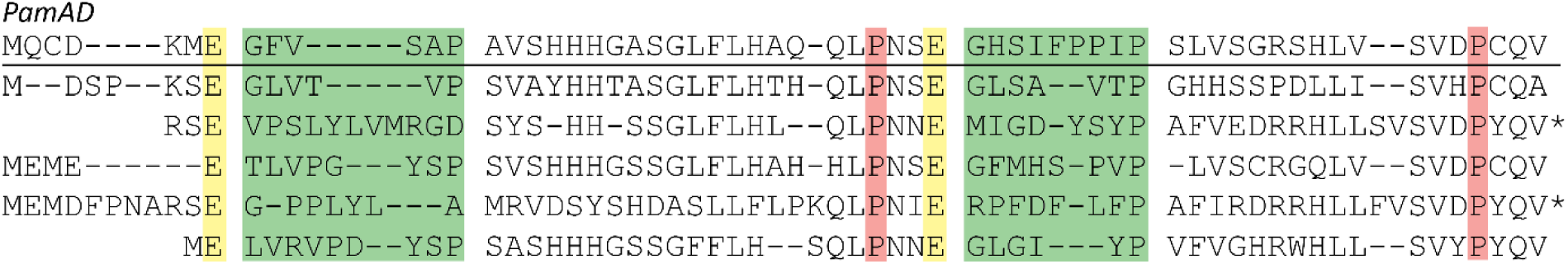
Manual alignment of the translated sequences of *Proannomuricatin* genes from the transcriptome of *A. muricata* seeds. The putative annomuricatin core peptide sequences are separated from the leader and follower sequences with spaces and have a green background. The absolutely conserved Glu in the P1 position is shown with a yellow background. Note the absolutely conserved Pro (pink background) which may be significant in processing of the peptide precursors, after cleavage at the *N*-terminus of each core peptide.

The annomuricatins described here are typical in size for orbitides at between six and nine residues. Many annomuricatins contain mainly hydrophobic residues such as Val, Gly, Ala, Leu and Ile, which again is typical of orbitides.^10^ Annomuricatin K is unusual in that it has an Arg and an Asp residue. Both residues are rare in most orbitides except for the PLPs, which have an absolutely conserved Asp at the *C*-terminus of the core peptide, essential for their macrocyclization.

### Other annomuricatin-like peptides

The number of similar annonomuricatins suggests the genes encoding them have duplicated and diverged. To investigate whether this type of gene is more widespread among the Annonaceae, RNA-seq data were downloaded from the NCBI Sequence Read Archive. These data were generated from RNA isolated from the mature flowers of *Annona squamosa*^30^ and leaves of *A. muricata*.^31^ We assembled transcriptomes and searched them with tBLASTn using the *pamAD* transcript as the query sequence. This approach identified several contigs with strong sequence similarity to *Pam* genes. Three candidate transcripts were identified in *A. muricata* leaves with a maximum expected value of 1.6×10^−3^, which were different from those in the *A. muricata* seeds, encoding a total of six putative peptides (Figure 6A). Ten candidate transcripts from *A. squamosa* (maximum expected value 2.7×10^−4^) were also found, most encoding two possible peptides, though several varied considerably from the canonical sequence, with the putative core peptide lacking a *C*-terminal Pro or Ala (Figure 6B).

**Figure 6.**
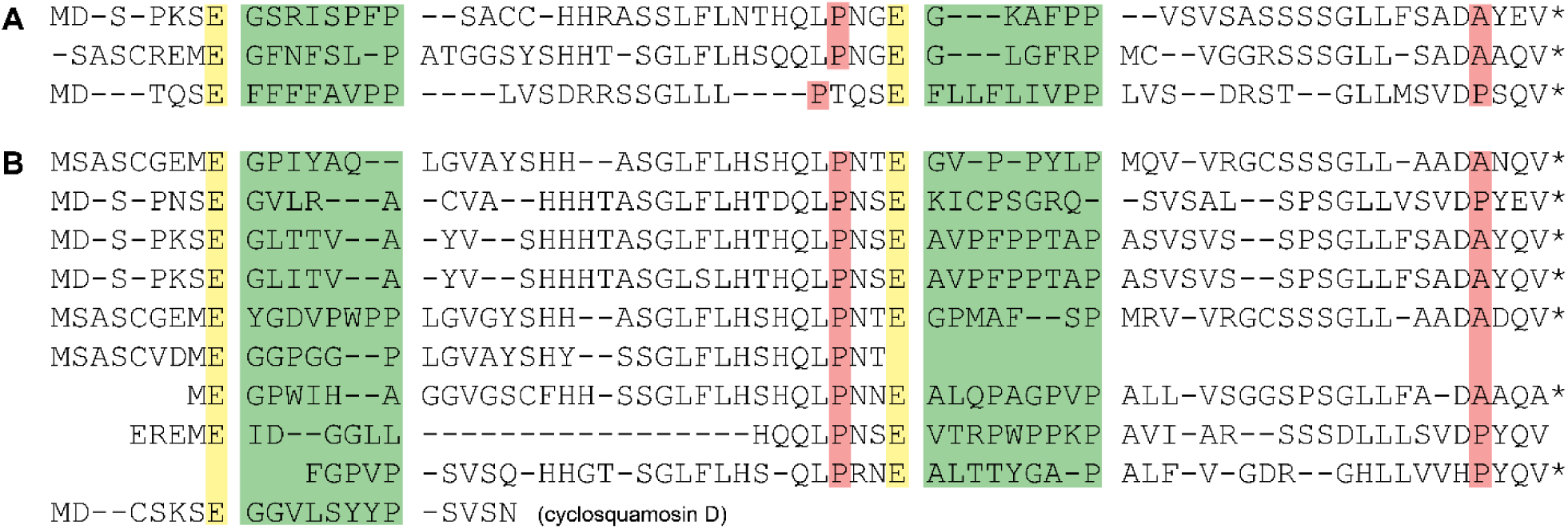
Alignments of transcripts from (A) *A. muricata* leaves and (B) *A. squamosa* flowers. Putative core peptides are shown with a green background, the absolutely conserved Glu with a yellow background, and the highly conserved Pro or Ala in pink. Asterisks are stop codons, shown where the ORF is complete. The last sequence shown corresponds to the known cyclic peptide, cyclosquamosin D,^32^ and appears to be only a fragment of the full ORF.

The cases where the putative core peptides do not have Pro or Ala at the *C*-terminus may represent genes where one of the two peptides encoded has degenerated into a non-functional sequence, as appears to have occurred in the *PamG* and *PamL* transcripts from *A. muricata* seeds. Again, the Glu in the P1 position at the *N*-terminus of the core peptide is absolutely conserved. The highly conserved Pro in the follower sequence is again prominent, though sometimes replaced by Ala. In one case it was in the fifth position from the *C*-terminus rather than the fourth. Without a tissue sample on which to perform MS/MS analysis, it was impossible to say whether these differences from the *A. muricata* seed sequences prevent production of the cyclic peptide.

What can be seen in these homologs of the *Pam* genes of *A. muricata* seeds is that similar transcripts are present in more than one Annonaceae species. The putative propeptides encoded by the transcripts have a high degree of sequence similarity and most of them appear to encode two cyclic peptides. We therefore suggest this type of orbitide-encoding gene could occur across the Annonaceae and would be an interesting topic for further research. It is also noteworthy that the putative peptides from the leaves of *A. muricata* are quite different to the seed peptides, suggesting the annomuricatins could have organ-specific functions.

### Biosynthesis of annomuricatins

The *Proannomuricatin* transcripts are short (∼200 nt) and the leader and follower sequences around the core peptide are highly conserved (Figure 5). The core peptide sequence is highly variable, but the *N*-terminus has a highly conserved Gly, preceded by an absolutely conserved Glu in the P1 position. The *C*-terminal Pro of the core peptide is also absolutely conserved, except for the presence of Ala in one instance.

Much work has been done on the biosynthesis of plant cyclic peptides that rely on AEP for their maturation and cyclisation.^4-8,33^ A similar depth of understanding exists for the orbitide segetalin A and its relatives, whose cyclization of a conserved *C*-terminal Ala residue is performed by the prolyl oligopeptidase PCY1.^14,29^ Prior to cyclization, the segetalins are cleaved at the *N*-terminus of the core peptide by an as-yet-uncharacterized enzyme, OLP1,^29^ which presumably is able to recognize the conserved Gln or Glu residue at the P1 position. Another example of peptides cyclized by a prolyl oligopeptidase are the amanitins from the fungus *Galerina marginata;* these are cyclized by GmPOPB. Unlike the segetalins, pro-amanitin is cleaved by POPB at the *N*-terminus of the core peptide as well as at the *C*-terminus due to the conserved Pro at the P1 position to the *N*-terminus.^34^ Based on the conserved residues in *Pam* genes and parallels with other cyclic peptide biosyntheses, annomuricatin precursors are likely to be cleaved first at the core peptide *N*-terminus by a Glu-targeting protease and then cleaved at the *C*-terminus and ligated by a POP. This may parallel the action of OLP1 and PCY1 in *V. hispanica*, since the former appears to target Glu or Gln at the proto-*N*-terminus, and the latter cleaves at Ala or Pro.

Here we have shown that one known and nine novel annomuricatins in the seeds of *A. muricata* are encoded by six very similar short genes; four of them encode two cyclic peptides, and the other two encode one peptide each. Similar genes are also found in the leaves of *A. muricata* and flowers of *A. squamosa*, indicating that such cyclic peptides may be present in other Annonaceae species. Comparison with the segetalins of *V. hispanica* indicates that a prolyl oligopeptidase is the likely cyclisation agent. Further study is required to identify this POP and to characterize its structure and mechanism of action.

## EXPERIMENTAL SECTION

### Plant material

Seeds of *A. muricata* were purchased from B & T World Seeds (Paguignan, France). While under quarantine, seeds were treated with Gaucho 600 insecticide (Bayer CropScience) to comply with Western Australian regulations. Soon after, seeds were frozen in liquid nitrogen, ground to a fine powder and stored at −80 °C until required.

### Seed peptide extraction

Seed peptides were extracted as previously described.^12^ Briefly, peptides were extracted in 50% MeOH / 50% CHCl_2_. Phases were separated by the alternate addition of CHCl_3_ and 0.05% trifluoroacetic acid. The upper, aqueous phase was dried overnight in a vacuum centrifuge (Labconco) prior to purification.

### Purification of seed peptide extracts

Seed peptide extracts were purified according to the method previously described.^13^ Briefly, the crude extract was purified by solid-phase extraction using a 30 mg Strata-X polymeric reversed-phase column (Phenomenex). The extract was applied to the column as an aqueous solution of 5% MeCN (v/v) / 0.1% formic acid (v/v), then purified peptides were eluted with 85% MeCN (v/v) / 0.1% formic acid (v/v). The extract was dried in a vacuum centrifuge and redissolved in 5% MeCN (v/v) / 0.1% formic acid (v/v) for LC-MS analysis. HPLC-grade solvents were used throughout (Honeywell).

### LC-MS/MS for peptide sequencing

Samples (2 µL) were injected onto an EASY-Spray PepMap C18 column (75 μm x 150 mm, 3 μm particle size, 10 nm pores; Thermo Fisher Scientific) using a Dionex UltiMate 3000 nano UHPLC system (Thermo Fisher Scientific) at flow rate of 200 nL/min by the “µL pick-up” method. A gradient elution was run from 5% solvent B to 95% solvent B over 40 minutes. Solvent A was 0.1% formic acid in water and solvent B was 0.1% formic acid in MeCN (Fisher Scientific). The resulting electrospray (source voltage 1,800 V) was analyzed by an Orbitrap Fusion mass spectrometer (Thermo Fisher Scientific) running in positive ionization, data-dependent, “top speed” MS/MS mode, employing the Orbitrap mass analyzer for both MS and MS/MS measurements at a resolution of 120,000 for MS and 60,000 for MS/MS. Parameters were set as follows: HCD fragmentation alternating between 14% ± 3% and 23% ± 3% energy, MS scan range from 400 to 1600 *m/z*, minimum MS/MS *m/z* 50, isolation window 1.2, ACG 400,000 (MS) and 500,000 (MS/MS), maximum injection time 200 ms (MS) and 250 ms (MS/MS) and 2 microscans for MS/MS. Only ions with an intensity > 100,000 were fragmented for MS/MS.

### Peptide sequencing

Peptides were sequenced by visual examination of MS/MS spectra, aided by fragment predictions from the program mMass^35^ for cyclic peptides and MS Product on the ProteinProspector website for acyclic peptides.^36^ The parameters used for MS-Product selected a, b, y and immonium ions, plus internal fragments. Neutral losses were set to water (when S, T, E or D present) or ammonia (when R, K, Q or N present). Other parameters were left at their default values. Similar options were chosen for mMass except that y ions were not selected; these do not appear in the mass spectra of cyclic peptides because such peptides lack a carboxyl-terminus.

### *Annona muricata* RNA-seq and transcriptome assembly

Seeds of *A. muricata* (100 mg) were ground to a powder using a mortar and pestle cooled with liquid nitrogen. Total RNA was isolated using the Spectrum Plant Total RNA kit (Sigma Aldrich) and quality validated on a TapeStation 2200 system (Agilent). RNA-seq libraries were generated using the TruSeq Stranded Total RNA with Ribo-Zero Plant kit according to the manufacturer’s instructions (Illumina) and sequenced on a NextSeq 550 system (Illumina) as paired-end reads with a length of150 bp and an average quality score (Q30) of above 90%. The raw reads were deposited in the NCBI Sequence Read Archive under accession number SRR8959862.

The *A. muricata* transcriptome was assembled using CLC Genomics Workbench 10.0.1 (QIAGEN Aarhus A/S). The raw reads were trimmed to a quality threshold of Q30 and minimum length 50, and the assembly was performed with word size 64 and minimum contig length 200. Other parameters remained at their default values.

### Other *Annona* transcriptome assemblies

RNA-seq paired-read data were downloaded from the Sequence Read Archive of the National Institutes of Health for *A. squamosa* mature flowers (run SRR3478571) and *A. muricata* leaves (run ERR2040135). Using CLC Genomics Workbench 11.0, transcriptomes were assembled for each of these datasets. For *A. squamosa*, the raw paired-end reads were trimmed to a quality threshold of 22 and minimum length 50, and the assembly was performed with word size 64, minimum contig length 50 and bubble size 100. For *A. muricata* leaves, the raw paired-end reads were trimmed to a quality threshold of 20 and minimum length 50, and the assembly was performed with word size 50 and minimum contig length 50. In both cases, all other parameters remained at their default values.

### BLAST searches of transcriptomic data

A BLAST database was constructed from each of the assembled transcriptomes using CLC Genomic Workbench. To search for proannomuricatin peptide sequences, the *A. muricata* database was searched using tBLASTn with the PamAD peptide sequence as a query. The substitution matrix used was PAM45 to find only closely related sequences. Other parameters selected were: maximum expected value 1, gap open cost 15, gap extend cost 2 and word size 3. Similar searches were carried out on transcriptomes of *A. muricata* leaves and *A. squamosa* flowers downloaded from the SRA.

### Cloning of *Proannomuricatin* genes

Genomic DNA was extracted from 2 g of frozen *A. muricata* seed powder with the DNEasy Mericon Food Kit (QIAGEN) according to the manufacturer’s instructions. DNA was quantified using a NanoDrop 2000 (Thermo Fisher Scientific).

The genomic DNA template was amplified by the polymerase chain reaction (PCR) using *Pfu* Ultra High-Fidelity DNA polymerase (Agilent Technologies). Each 50 μL reaction consisted of genomic DNA (∼12 ng), 5 μL *Pfu* Ultra DNA polymerase reaction buffer (10x), 400 μM mixed dNTPs, 0.5 μL *Pfu* Ultra DNA polymerase and 0.4 μM of each of the appropriate forward and reverse primers (Table S1). PCR amplification was performed in a Veriti 96-well thermocycler (Applied Biosystems) programmed as follows: 95 °C for 2 min followed by 5 cycles of 95 °C for 30 s; 65 °C for 30 s; 72 °C for 30 s then 30 cycles of 95 °C for 30 s; 60 °C for 30 s; 72 °C for 30 s; and finally 72 °C for 10 min (*PamAD, PamEF, PamG*), or 95 °C for 2 min followed by 35 cycles of 95 °C for 30 s; 60 °C for 30 s; 72 °C for 30 s; and finally 72 °C for 10 min (*PamHI, PamJK, PamLM*).

PCR products were purified using the QIAquick PCR Purification Kit (QIAGEN) according to the manufacturer’s instructions. DNA was eluted in 30 µL of water, quantified on a NanoDrop 2000 and sent for dideoxy sequencing using the forward PCR primers mentioned above (Garvan Institute, Darlinghurst NSW, Australia).

The six *Proannomuricatin* (*Pam*) genes from this study were deposited in GenBank under accession numbers MK836460-MK836465.

## AUTHOR INFORMATION

### Author Contributions

M.F.F. and J.S.M. conceived the study; O.B. and J.W. performed RNA-seq; J.Z. and M.F.F. assembled the *A. muricata* transcriptome; M.F.F performed all other experiments and analyzed data; M.F.F and J.S.M. wrote the manuscript with help from all other authors.

### Notes

The authors declare no competing financial interest.

## ACKNOWLEDGMENTS

The authors thank Nicolas L. Taylor of the School of Molecular Sciences at the University of Western Australia for providing advice and assistance in mass spectrometry aspects of this project.

The authors acknowledge the facilities, and the scientific and technical assistance of the Australian Microscopy & Microanalysis Research Facility at the Centre for Microscopy, Characterisation & Analysis, The University of Western Australia, a facility funded by the University, State and Commonwealth governments.

## SUPPORTING INFORMATION

### Supplementary Tables

**Table S1.**
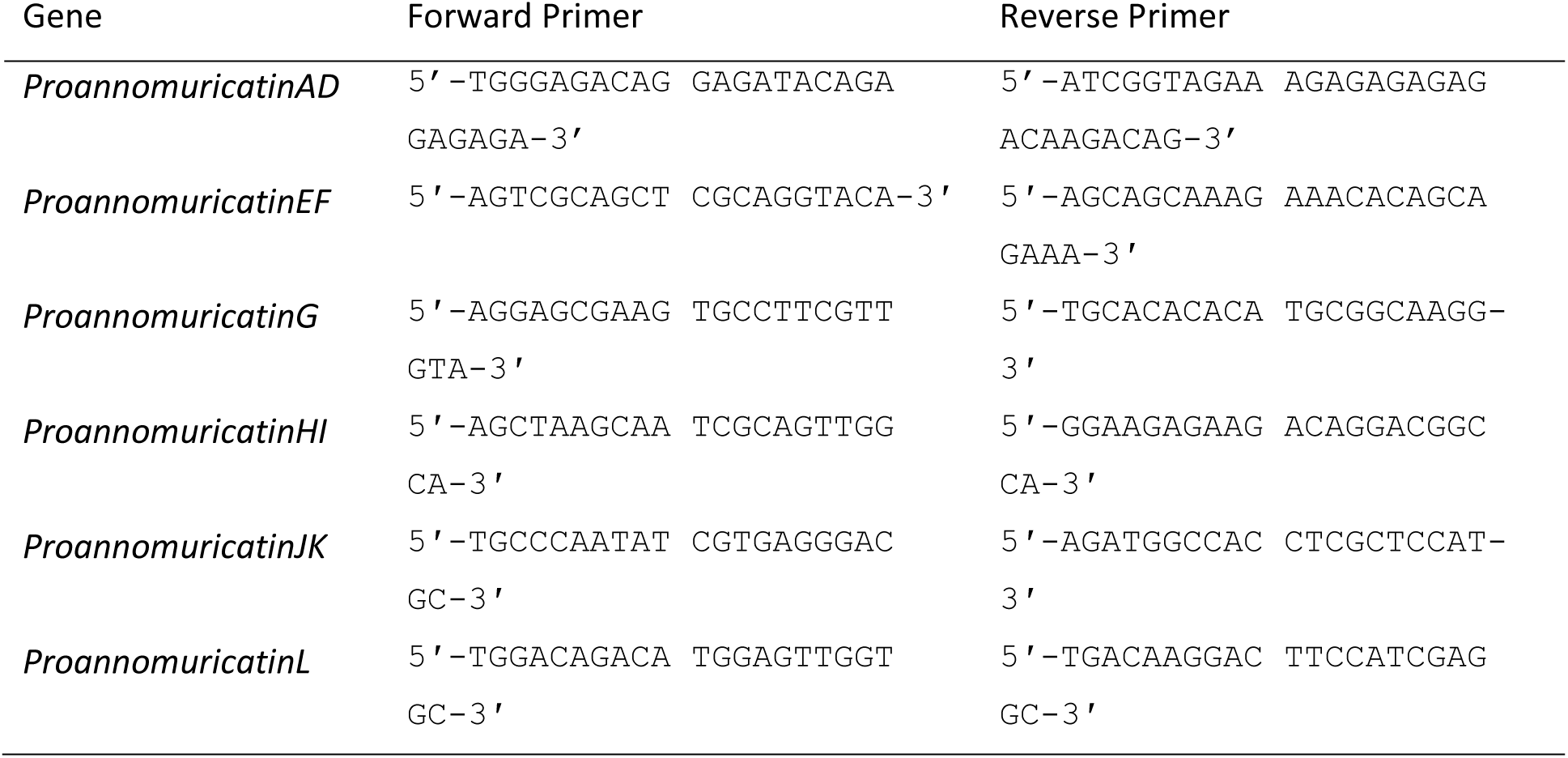
PCR primers used for the amplification and sequencing of the putative *Proannomuricatin* genes identified in transcriptome data.

### Supplementary Figures

**Figure S1.**
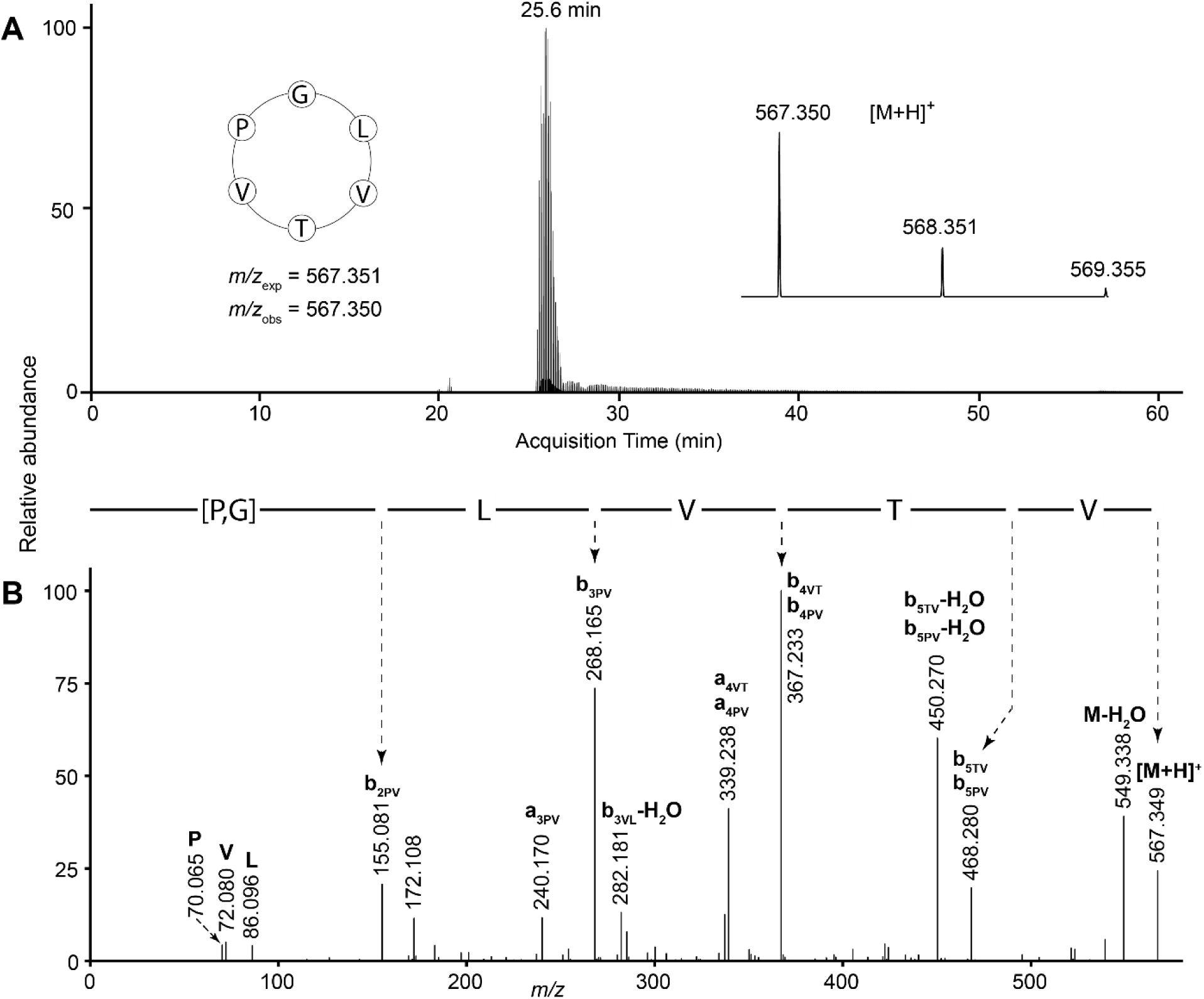
LC-MS data for annomuricatin E with sequence cyclo-GLVTVP. (A) Extracted ion chromatogram showing acquisition time of the peptide, with (inset left) peptide sequence with expected and observed mass-to-charge ratios (*m/z*) and (inset right) peptide mass spectrum. (B) Tandem mass spectrum of the fragmented precursor ion. Immonium ions are denoted by the one-letter code of the residue they represent. The derived amino acid sequence is shown above the spectrum; residues in square brackets could not be assigned an order.

**Figure S2.**
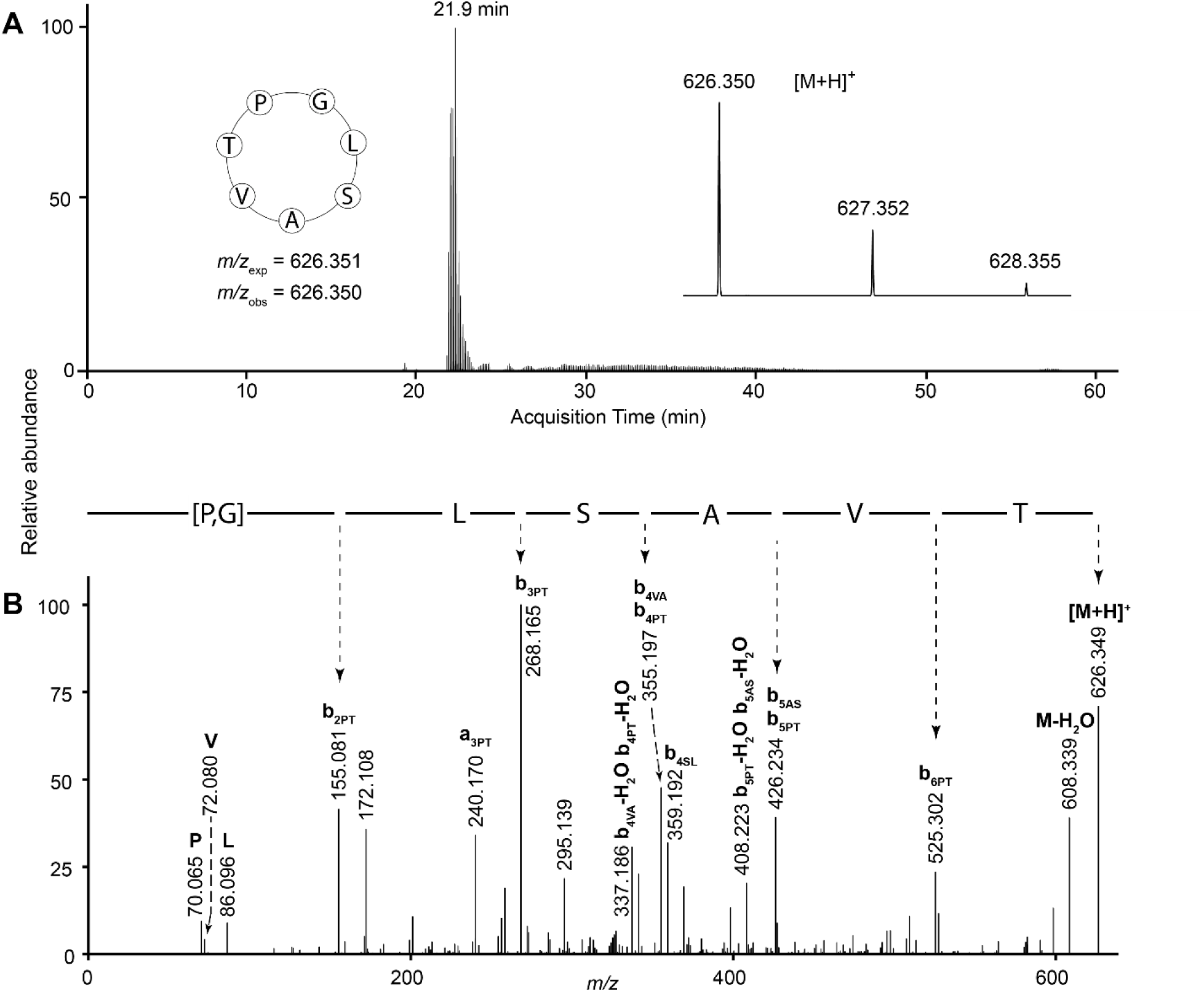
LC-MS data for annomuricatin F with sequence cyclo-GLSAVTP. (A) Extracted ion chromatogram showing acquisition time of the peptide, with (inset left) peptide sequence with expected and observed mass-to-charge ratios (*m/z*) and (inset right) peptide mass spectrum. (B) Tandem mass spectrum of the fragmented precursor ion. Immonium ions are denoted by the one-letter code of the residue they represent. The derived amino acid sequence is shown above the spectrum; residues in square brackets could not be assigned an order.

**Figure S3.**
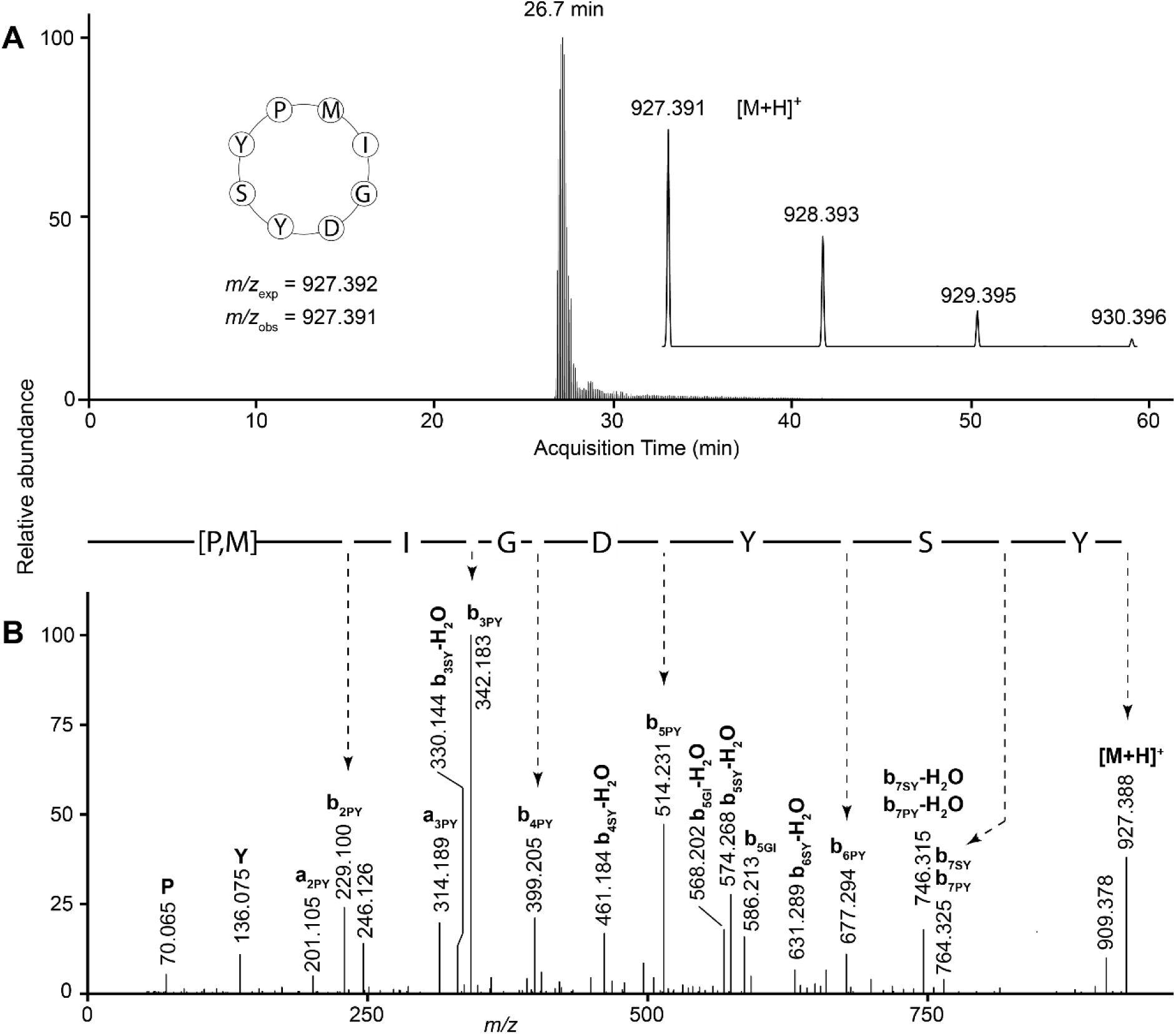
LC-MS data for annomuricatin G with sequence cyclo-MIGDYSYP.(A) Extracted ion chromatogram showing acquisition time of the peptide, with (inset left) peptide sequence with expected and observed mass-to-charge ratios (*m/z*) and (inset right) peptide mass spectrum. (B) Tandem mass spectrum of the fragmented precursor ion. Immonium ions are denoted by the one-letter code of the residue they represent. The derived amino acid sequence is shown above the spectrum; residues in square brackets could not be assigned an order.

**Figure S4.**
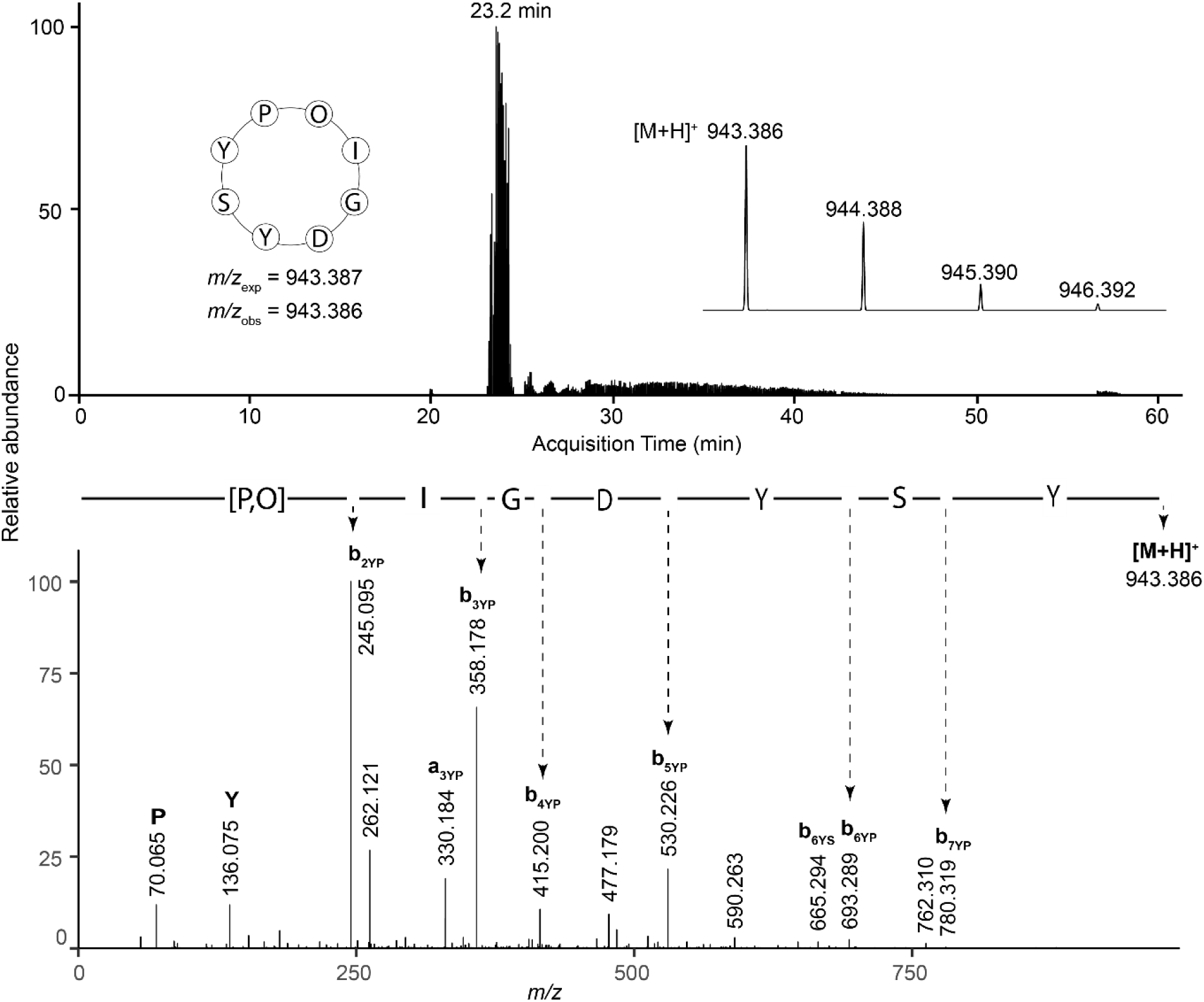
LC-MS data for annomuricatin G with sequence cyclo-MIGDYSYP, although the Met was oxidized (hence O in figure) (A) Extracted ion chromatogram showing acquisition time of the peptide, with (inset left) peptide sequence with expected and observed mass-to-charge ratios (*m/z*) and (inset right) peptide mass spectrum. (B) Tandem mass spectrum of the fragmented precursor ion. Immonium ions are denoted by the one-letter code of the residue they represent. The derived amino acid sequence is shown above the spectrum; residues in square brackets could not be assigned an order.

**Figure S5.**
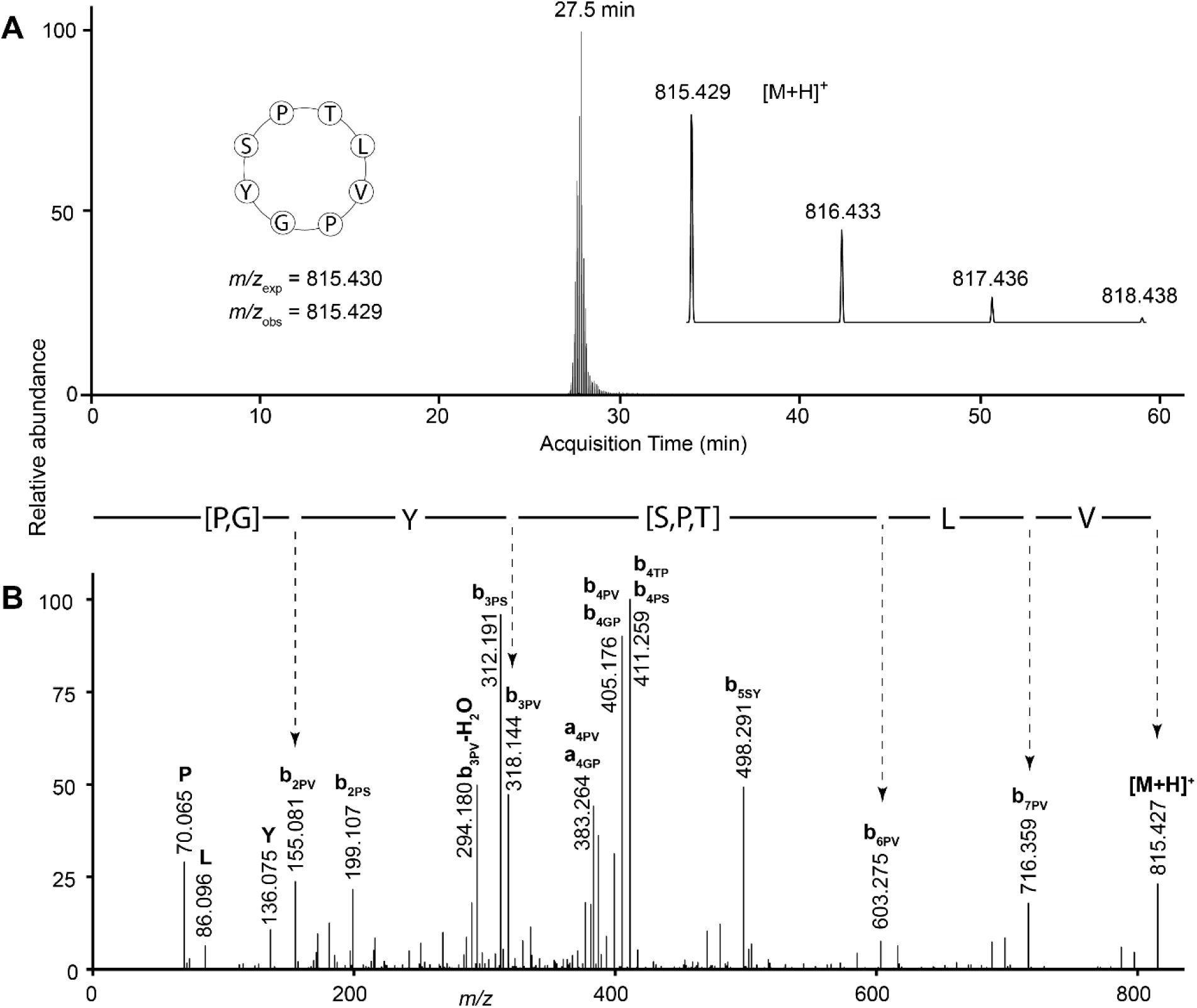
LC-MS data for annomuricatin H with sequence cyclo-TLVPGYSP. (A) Extracted ion chromatogram showing acquisition time of the peptide, with (inset left) peptide sequence with expected and observed mass-to-charge ratios (*m/z*) and (inset right) peptide mass spectrum. (B) Tandem mass spectrum of the fragmented precursor ion. Immonium ions are denoted by the one-letter code of the residue they represent. The derived amino acid sequence is shown above the spectrum; residues in square brackets could not be assigned an order.

**Figure S6.**
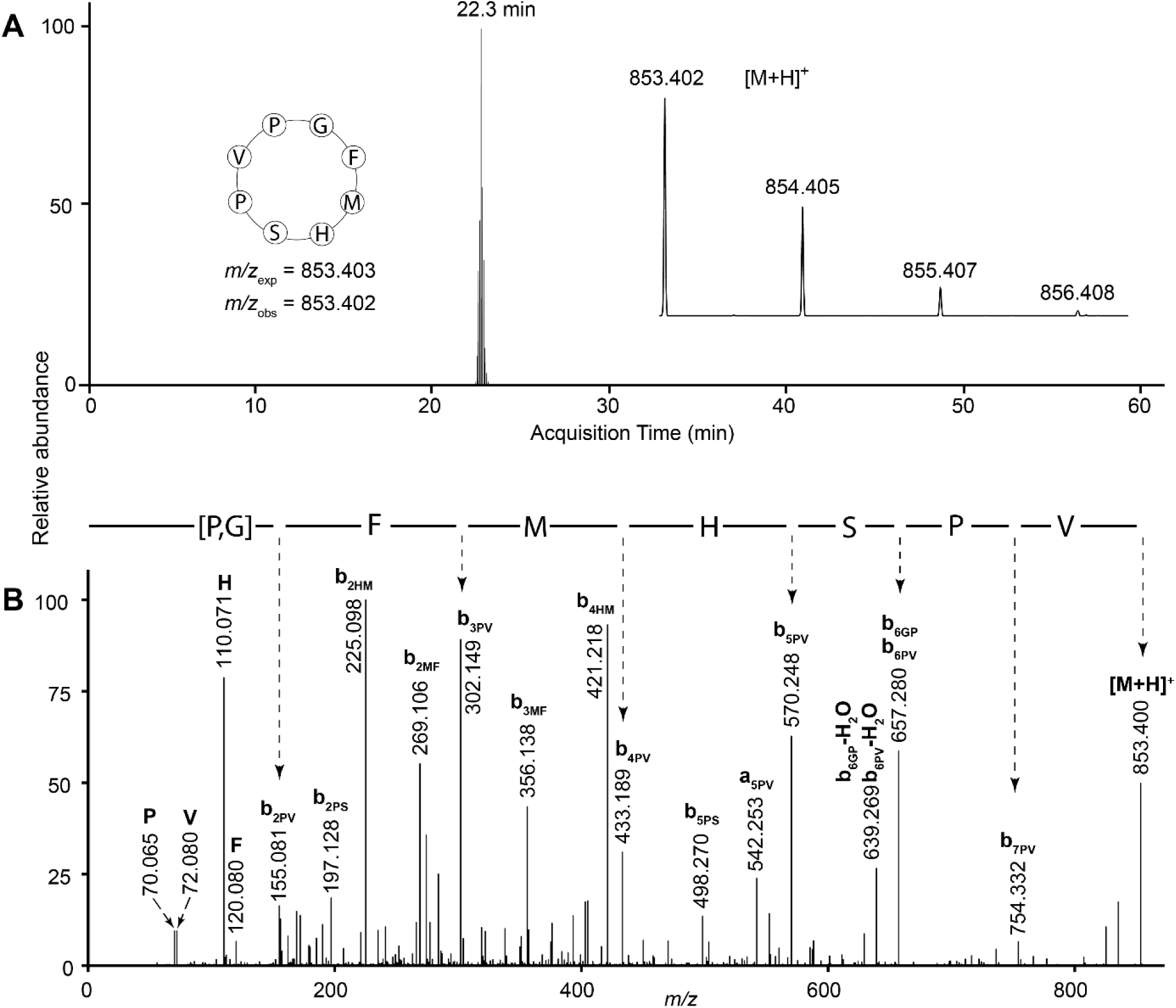
LC-MS data for annomuricatin I with sequence cyclo-GFMHSPVP. (A) Extracted ion chromatogram showing acquisition time of the peptide, with (inset left) peptide sequence with expected and observed mass-to-charge ratios (*m/z*) and (inset right) peptide mass spectrum. (B) Tandem mass spectrum of the fragmented precursor ion. Immonium ions are denoted by the one-letter code of the residue they represent. The derived amino acid sequence is shown above the spectrum; residues in square brackets could not be assigned an order.

**Figure S7.**
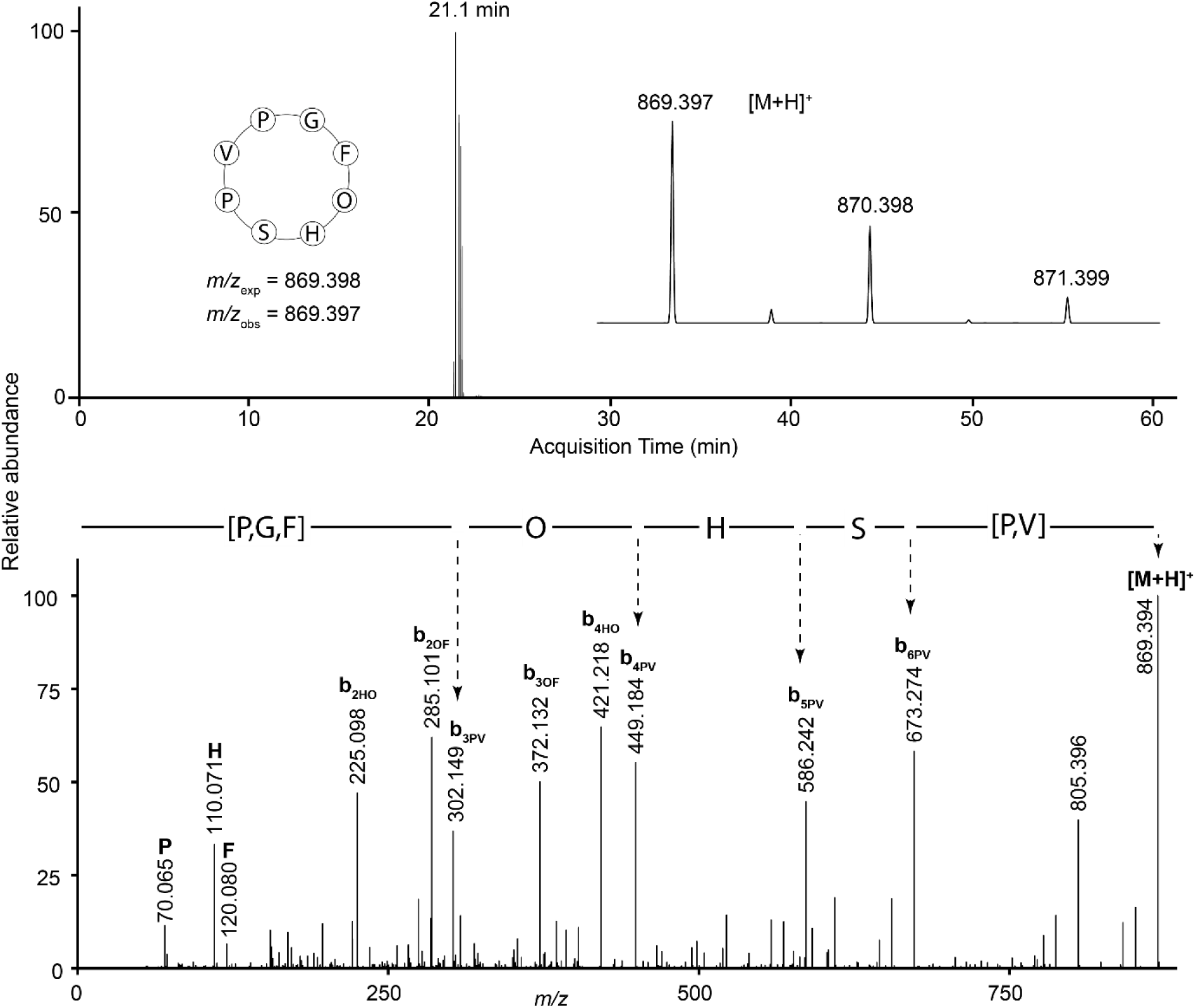
LC-MS data for annomuricatin I with sequence cyclo-GFMHSPVP, although the Met was oxidized (hence O in figure).(A) Extracted ion chromatogram showing acquisition time of the peptide, with (inset left) peptide sequence with expected and observed mass-to-charge ratios (*m/z*) and (inset right) peptide mass spectrum.(B) Tandem mass spectrum of the fragmented precursor ion. Immonium ions are denoted by the one-letter code of the residue they represent. The derived amino acid sequence is shown above the spectrum; residues in square brackets could not be assigned an order.

**Figure S8.**
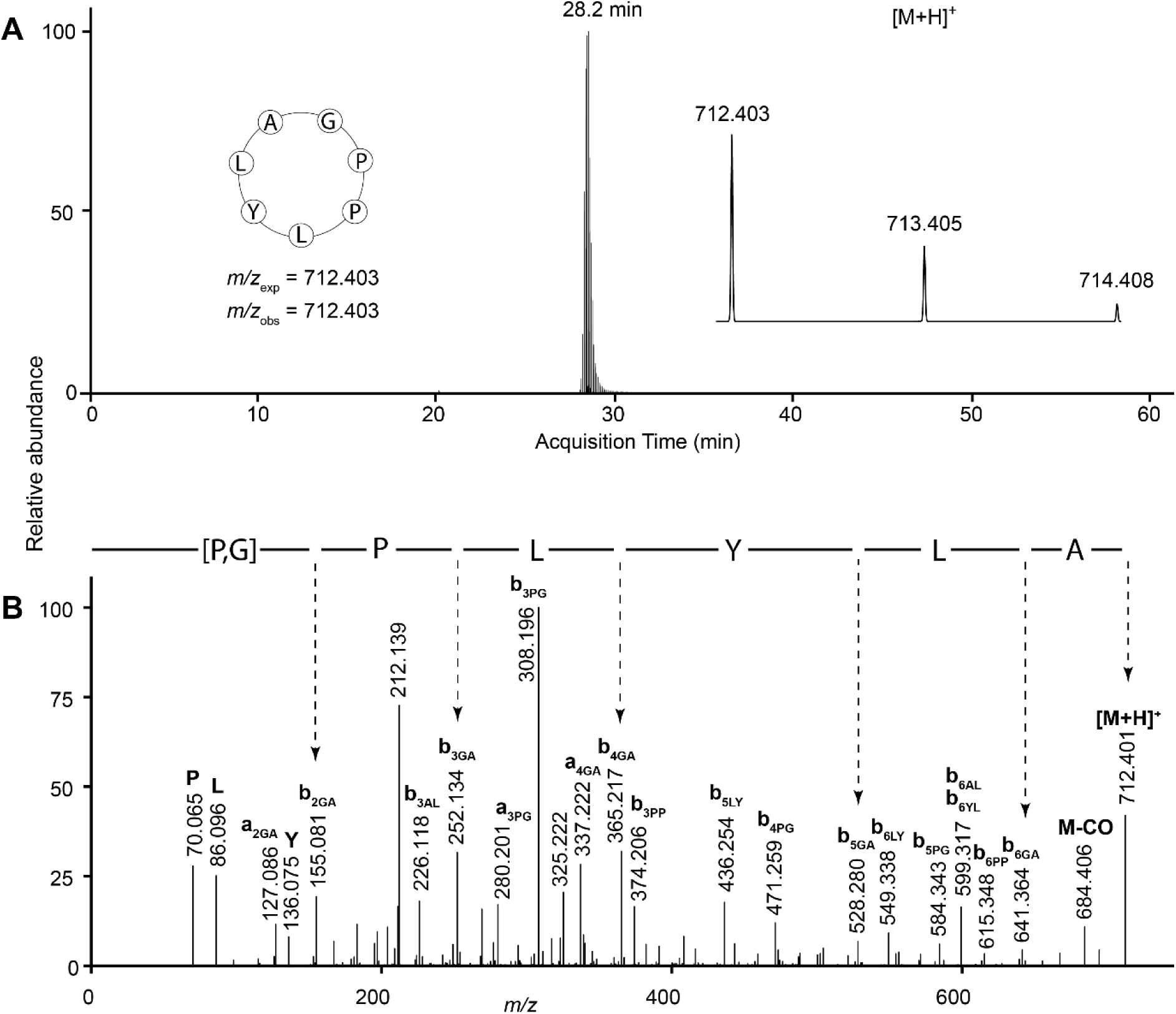
LC-MS data for annomuricatin J with sequence cyclo-GPPLYLA.(A) Extracted ion chromatogram showing acquisition time of the peptide, with (inset left) peptide sequence with expected and observed mass-to-charge ratios (*m/z*) and (inset right) peptide mass spectrum. (B) Tandem mass spectrum of the fragmented precursor ion. Immonium ions are denoted by the one-letter code of the residue they represent. The derived amino acid sequence is shown above the spectrum; residues in square brackets could not be assigned an order.

**Figure S9.**
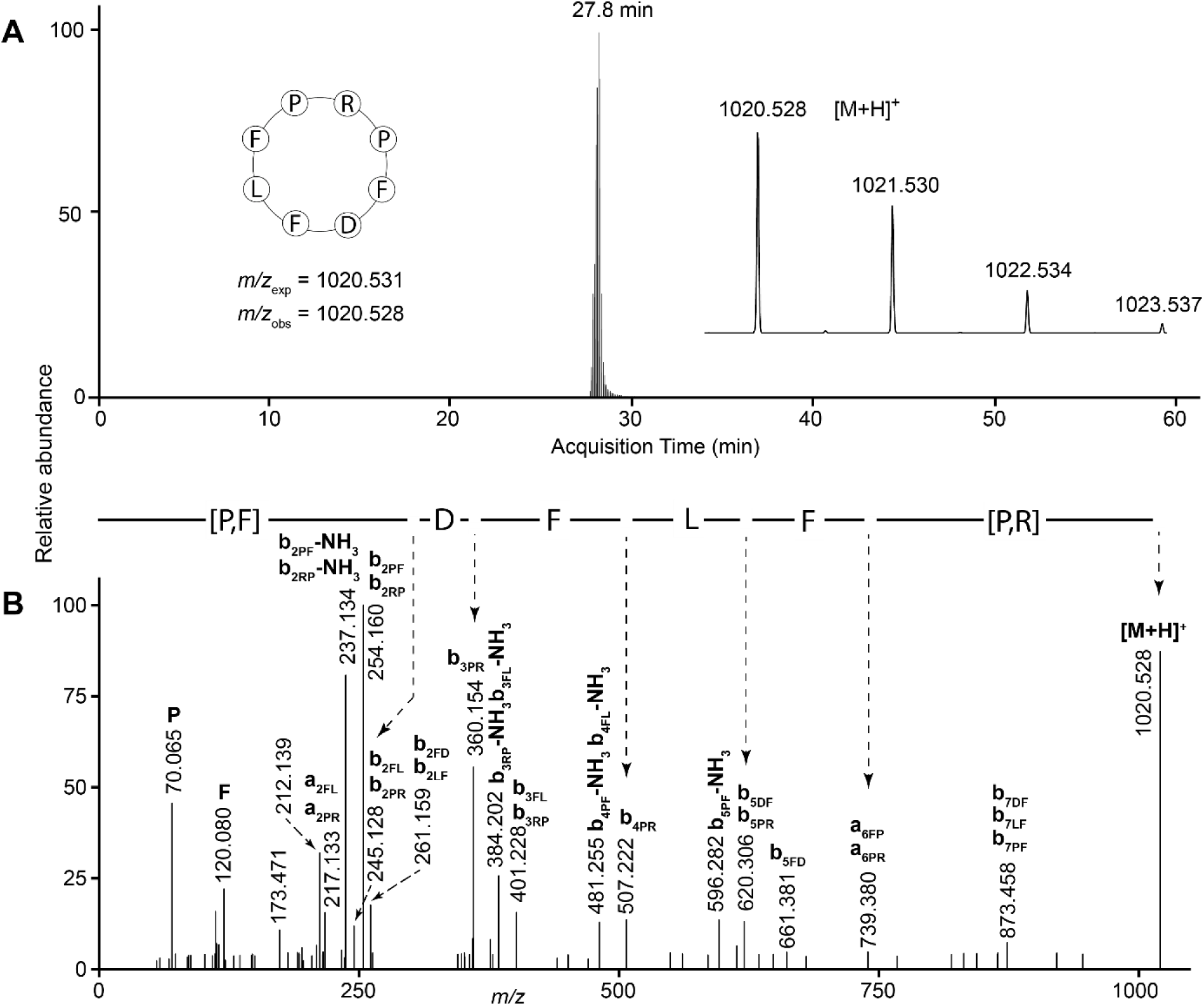
LC-MS data for annomuricatin K with sequence cyclo-RPFDFLFP. (A) Extracted ion chromatogram showing acquisition time of the peptide, with (inset left) peptide sequence with expected and observed mass-to-charge ratios (*m/z*) and (inset right) peptide mass spectrum. (B) Tandem mass spectrum of the fragmented precursor ion. Immonium ions are denoted by the one-letter code of the residue they represent. The derived amino acid sequence is shown above the spectrum; residues in square brackets could not be assigned an order.

**Figure S10.**
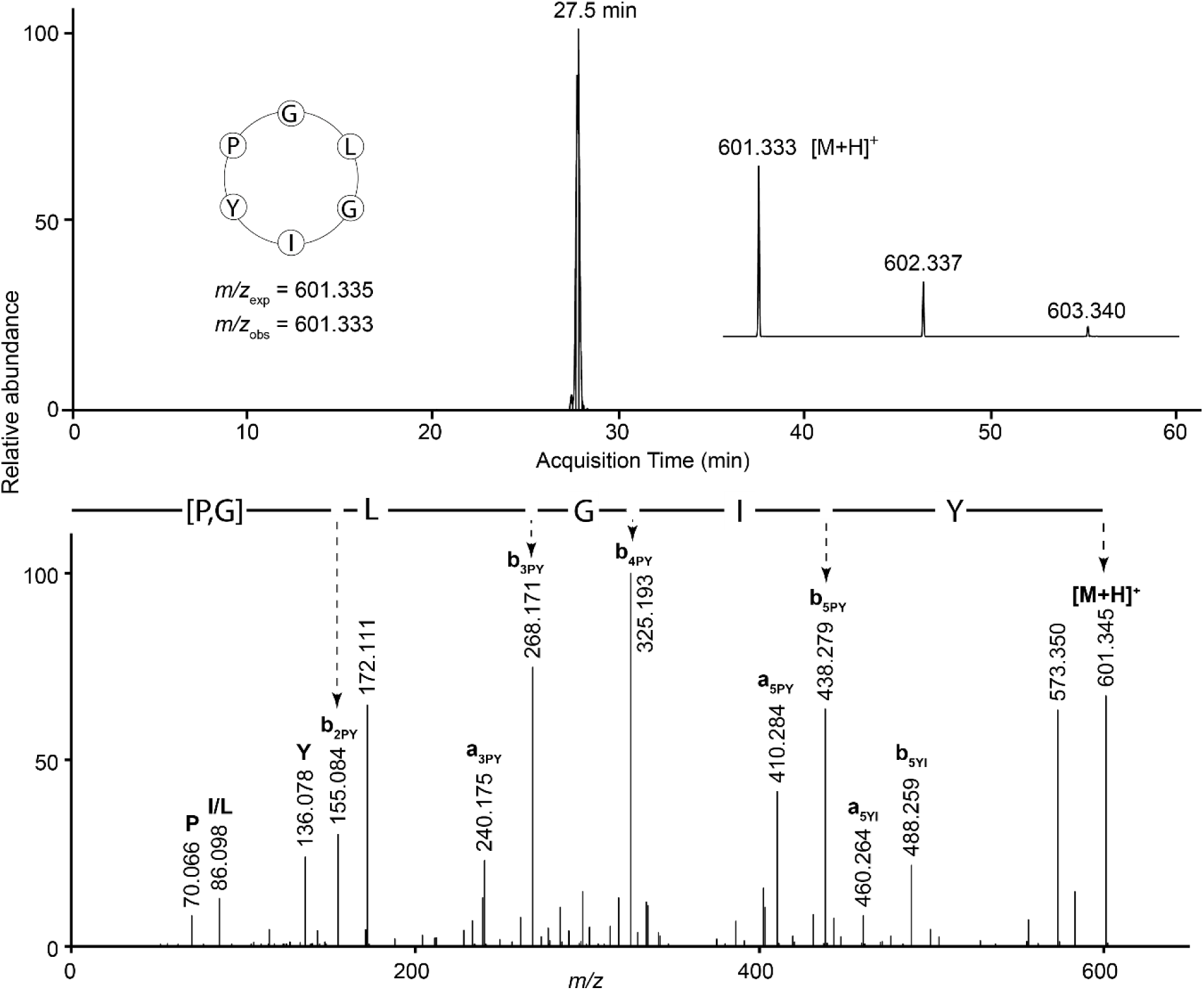
LC-MS data for annomuricatin L with sequence cyclo-GLGIYP. (A) Extracted ion chromatogram showing acquisition time of the peptide, with (inset left) peptide sequence with expected and observed mass-to-charge ratios (*m/z*) and (inset right) peptide mass spectrum. (B) Tandem mass spectrum of the fragmented precursor ion. Immonium ions are denoted by the one-letter code of the residue they represent. The derived amino acid sequence is shown above the spectrum; residues in square brackets could not be assigned an order.

**Figure S11.**
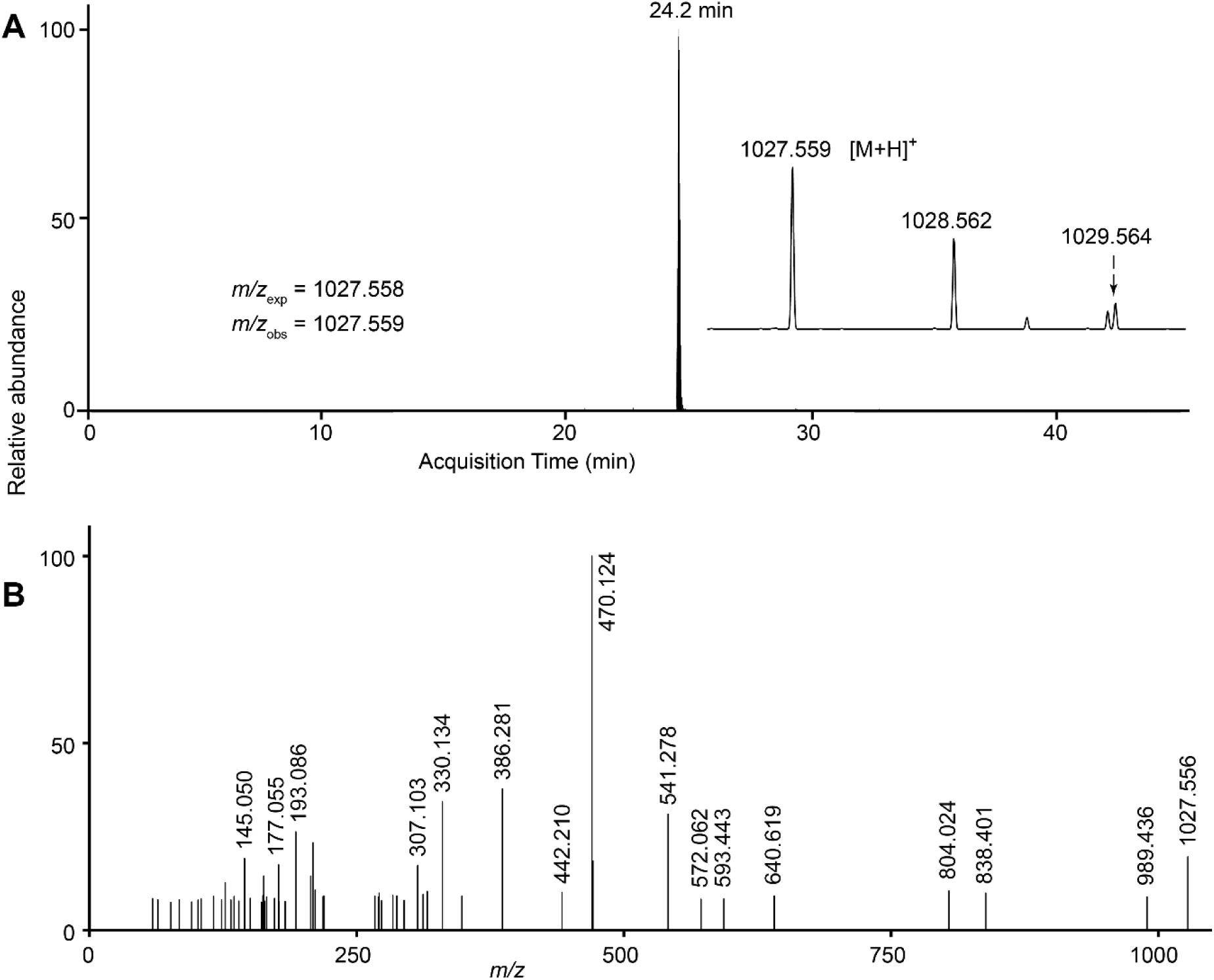
LC-MS data for putative peptide at *m/z* 1027.558. (A) Extracted ion chromatogram showing acquisition time of the peak, with (inset left) expected and observed mass-to-charge ratios (*m/z*) and (inset right) mass spectrum. **(B)** Tandem mass spectrum of the fragmented precursor ion.

### TABLE OF CONTENTS/ABSTRACT GRAPHIC

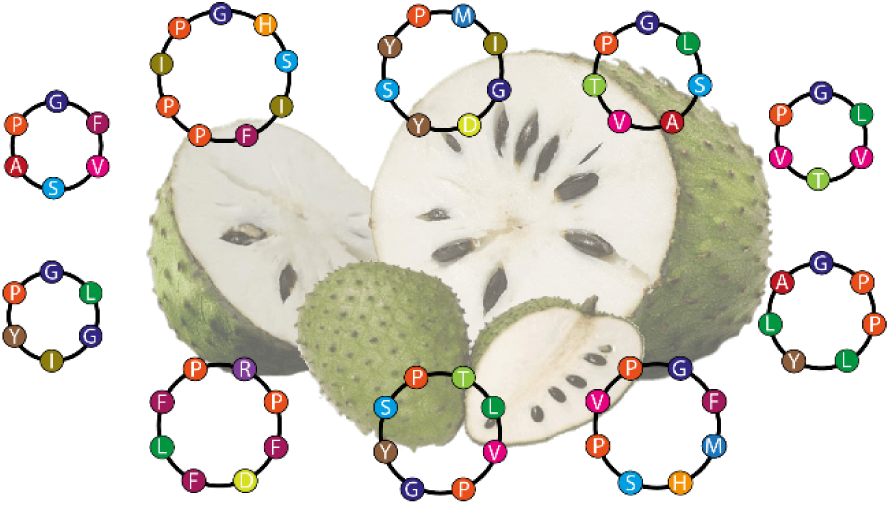

